# Resynchronization of the biological clock using exposure to low oxygen levels in humans: an exploratory study

**DOI:** 10.1101/2025.09.09.674331

**Authors:** Renée Morin, Geneviève Forest, Kim Isabelle-Nolet, Vincent Bourgon, Félix-Gabriel Duval, Jean-François Mauger, Pascal Imbeault

**Affiliations:** Behavioural and Metabolic Research Unit, University of Ottawa, Ottawa, ON, Canada; Université du Québec en Outaouais, Gatineau, Québec, Canada; Institut du Savoir Montfort, Montfort Hospital, Ottawa, ON, Canada

## Abstract

**Introduction:** Circadian desynchronization, evident in scenarios such as jet lag, shift work, and circadian rhythm sleep disorders, detrimentally affects sleep quality and overall health, both acutely and chronically. Aligning the circadian system with environmental time cues is therefore essential for maintaining physiological and psychological well-being. While luminotherapy and melatonin supplementation are widely used to facilitate circadian realignment, emerging evidence suggests that fluctuations in oxygen levels may also help in circadian realignment. However, the effect of hypoxia on the synchronisation of the circadian clock in humans remains largely unexplored.

**Methods:** Using a randomized controlled crossover study design, 11 healthy participants (6 men, 5 women, mean age 23.3 ± 1.9 years) completed one baseline condition and two 48-h experimental conditions. The baseline condition (BL) was used to establish circadian markers. Both experimental conditions simulated a phase advance of 4 hours. In condition 1 (Hypo), participants underwent a 2-hour normobaric hypoxic exposure (FiO_2_ = 12%), starting 2 h after habitual wake time. In condition 2 (Lum+Mel), participants received a 3-hour luminotherapy session (500 nm, 506 lux) at the same time point, combined with 5 mg of exogenous melatonin administered 6 hours before usual bedtime. Salivary melatonin levels were measured in each phase of the study to assess circadian phase shifts. Data were analyzed using linear mixed models.

**Results:** Salivary melatonin levels increased progressively over time in all conditions (p < 0.001), with significant differences observed between experimental conditions (p < 0.001), but no interaction effect (p = 0.854). Exposure to hypoxia significantly reduced oxyhemoglobin saturation (p < 0.05) and increased heart rate and subjective symptoms of fatigue. In terms of circadian phase, the dim light melatonin onset (DLMO) occurred 1.30 hour (78 minutes) earlier in the Lum+Mel condition compared to baseline (p=0.001). In the hypoxia condition, the DLMO occurred on average 0.58 hour (34.8 minutes) earlier than baseline, but this change did not reach statistical significance (p=0.156).

**Conclusions:** This study provides preliminary evidence that normobaric hypoxia may modestly advance the human circadian phase, although not to a statistically significant extent. In contrast, combined phototherapy and melatonin administration produced a robust and significant phase advance in salivary melatonin onset. These findings suggest that while hypoxia may influence circadian timing, established interventions like light and melatonin remain more effective for circadian realignment. Further research is warranted to elucidate the mechanisms and optimize the application of hypoxia in circadian modulation.

## Introduction

With the earth’s rotation creating a roughly 24-hour cycle of light and temperature, most life on this planet has evolved with internal biological clocks that anticipate and adapt to these daily shifts (Kuhlman et al., 2018). These circadian clocks synchronize our physiological processes to our environment, influencing sleep-wake cycles, body temperature, and hormone levels through cues known as *zeitgebers*—such as light exposure, eating patterns, and social activities (Mistlberger & Rusak, 2005). In humans, this synchronization is crucial for optimal functioning, as disruptions in circadian timing have been linked to increased risks of mental, metabolic, and cardiovascular disease, along with diminished alertness and workplace performance (Lewis et al., 2018; Paul et al., 2011). However, modern lifestyles increasingly challenge this synchronization. Global travel and shift work, for example, contribute to chronic circadian misalignment, manifesting as jet lag and irregular sleep patterns (Reid & Abbott, 2015). Under typical conditions, circadian rhythms are entrained daily to the 24-hour day/night cycle, with light serving as the primary synchronizer and melatonin playing a central role in regulating circadian timing. Consequently, interventions such as luminotherapy (i.e., light therapy) and exogenous melatonin administration have been widely studied and applied to facilitate circadian resynchronization (Cheng et al., 2021). Combined light therapy and exogenous melatonin are significantly more effective than individual treatments for advancing the circadian phase, though their effects are additive rather than synergistic (Burke et al., 2013; Cheng et al., 2021; Crowley & Eastman, 2012). The efficacy of these methods, however, varies depending on factors such as timing, intensity, dosage, and duration of exposure and individual differences (Bin et al., 2019; Cheng et al., 2021) underscoring the need for alternative or complementary strategies.

To assess the effectiveness of such interventions, researchers often rely on specific physiological markers of circadian timing. One key marker is the Dim Light Melatonin Onset (DLMO), which refers to the time at which melatonin levels first begin to rise under dim light conditions. It is considered the gold standard for assessing the phase of the central circadian clock (Hofstra & de Weerd, 2008; Weinert & Waterhouse, 2017). Compared to other circadian markers, the melatonin rhythm is relatively stable and less susceptible to external influences, making it a reliable indicator of circadian timing (Molina & and Burgess, 2011). However, a significant limitation to its widespread use in research is the high cost of melatonin assays. Nevertheless, the effectiveness of luminotherapy and exogenous melatonin, whether used alone or in combination, in shifting the circadian phase of DLMO is well documented (Cheng et al., 2021). For example, Burke and colleagues (Burke et al., 2013) demonstrated that a single 3-hour session of light exposure at 3000 lux, starting one hour before the usual wake time, led to a 0.70-hour DLMO phase advance. Similarly, administering 5 mg of exogenous melatonin 5.75 hours before bedtime resulted in a 0.62-hour advance. When combined, the two treatments resulted in a cumulative DLMO advance of 1.13 hours (Burke et al., 2013). Paul and colleagues (Paul et al., 2011) reported comparable findings: one hour of morning bright light (350 lux, green light) caused a non-significant DLMO phase advance of 0.31 hours, while a single 3 mg dose of exogenous melatonin taken at 4:00 PM led to a 0.72-hour advance. Their combination resulted in a 1.04-hour phase advance.

Given these findings, light therapy and exogenous melatonin have demonstrated measurable effects on circadian phase even after a single administration. However, despite their proven efficacy under controlled conditions, these interventions face several practical limitations that hinder their broader implementation. Current guidelines often lack consensus on optimal dosing and precise timing, with recommended administration windows that can be difficult to follow in everyday life—particularly for individuals facing abrupt schedule shifts, such as during transmeridian travel or shift work. As a result, while these strategies are widely recommended, their application in real-world settings remains constrained by the challenges of personalization, adherence, and accessibility. This highlights the need to either refine current protocols to enhance feasibility or explore alternative strategies that might offer a more robust and convenient “one-shot” solution for circadian resynchronization. In this context, we propose investigating hypoxemia (i.e., reduced blood oxygen levels) as a potential tool to induce voluntary circadian phase shifts. While not yet widely studied as a chronotherapeutic intervention, this approach is grounded in recent evidence—such as the work by Adamovich and colleagues (Adamovich et al., 2017)—suggesting that oxygen availability may influence circadian timing mechanisms.

Indeed, Adamovich and colleagues demonstrated that: 1) O_2_ levels display daily rhythms in rodent blood and tissues, which influence circadian clock genes in a hypoxia-inducible factor 1 (HIF1α)–dependent manner, and 2) adjustments in O_2_ levels were found to accelerate the adaptation in wild-type mice subjected to simulated jet lag. In their study, (6-hour phase advance), mice exposed to 12 hours of moderate hypoxia (16% O_2_) prior to the shift adapted significantly faster than control mice kept at normoxic levels (21% O_2_). Remarkably, even a short, 2-hour hypoxic pulse at 14% O_2_ administered after the phase shift led to a faster adjustment compared to controls — highlighting not only the sensitivity of the clock to O_2_ levels but also the potential of brief hypoxic exposures to enhance circadian realignment (Adamovich et al., 2017).

These findings suggest that hypoxia may offer a novel mechanism for circadian phase adjustment. While hypoxia has been associated with changes in melatonin levels and core body temperature rhythms (Coste et al., 2009), evidence for its direct effect on human circadian phase shifting remains limited. One study exposing participants to mild hypobaric hypoxia—simulating altitudes of 8,000 to 12,000 feet—for 8 hours (08:00 to 16:00) found that circadian markers such as core body temperature and endogenous melatonin were altered (phase delay) for more than 24 hours post-exposure (Coste et al., 2007). Earlier studies by the same group suggest that daytime hypoxia may induce a phase delay (Coste, Beaumont, et al., 2004; Coste et al., 2004). Though, only one time point was tested. Determining whether hypoxia follows a CRP is crucial for understanding its potential as a phase-shifting intervention. While both rodents and humans rely on HIF1α for oxygen homeostasis, differences in diurnal vs. nocturnal oxygen consumption rhythms complicate cross-species extrapolation. Importantly, we still lack data on whether hypoxia directly influences the central clock. More recently, a study by Post and colleagues showed that an exposure to a 6.5-hour interval of normobaric hypoxia (FiO_2_ ∼15%) early in the night in healthy adults advanced melatonin onset by 9 minutes following a constant posture protocol. These results provide direct evidence that hypoxia can act as a circadian time cue in humans, potentially influencing clock timing through the body’s oxygen-sensing pathways (Post et al., 2025).

To our knowledge, only few studies have demonstrated hypoxia-induced circadian phase shifts under controlled conditions (Coste, Beaumont, et al., 2004; Coste et al., 2004, 2009; Post et al., 2025). Accordingly, this exploratory study aimed to investigate whether exposure to reduced oxygen levels can facilitate circadian resynchronization in humans undergoing a phase advance protocol. Advancing the circadian phase refers to a forward shift in the timing of the endogenous biological rhythms, resulting in earlier onset of physiological processes such as melatonin secretion, sleep initiation, and core body temperature nadir. More precisely, we investigated whether acute normobaric hypoxia can induce phase shifts in human circadian rhythms comparable to those produced by a combined intervention of light therapy and exogenous melatonin during a phase advance protocol. Based on prior findings, we hypothesized that hypoxic exposure would promote a phase advance in the circadian rhythm of salivary melatonin in response to a simulated 4-hour jet lag scenario comparatively to a well-known intervention such as luminotherapy combined with melatonin.

## Methodology

### Participants

Fifteen healthy participants, eight men and seven women, ages 18-35, were recruited for this exploratory study. Four participants withdrew from the study at various stages of the research protocol. Of these individuals, two cited scheduling conflicts, while another reported difficulty sleeping in the laboratory and experienced migraines during the week following the hypoxic condition. The final individual withdrew due to health issues unrelated to the research project. Therefore, the final sample size was eleven participants. The participants were recruited using posters installed at the University of Ottawa (UO) and at the Université du Québec en Outaouais (UQO), through social media platforms (e.g., Facebook). All participants provided written informed consent prior to participation. The study protocol received ethical approval from the Research Ethics Board of both UO and UQO. To minimize potential physiological or behavioral confounds, inclusion and exclusion criteria were applied. Exclusion criteria included: presence of sleep, neurological, psychiatric, or medical disorders, history of depression, anxiety or diabetes, use of prescription medications known to affect sleep, habitual napping, habitual sleep duration less than 7 hours or more than 9 hours per night,; shift work within the past year, use of illicit drugs, nicotine or excessive consumption of caffeine or alcohol, and a BMI outside the range of 18.5 to 29.9. Inclusion criteria required participants to maintain a consistent sleep schedule, with bedtimes between 10 p.m. and midnight and wake times between 6 a.m. and 9 a.m., and to regularly obtain 7-9 hours of sleep per night. Female participants were required to use monophasic oral contraceptive to control for hormonal fluctuations that could influence sleep and physiological measures. Participants who had recently traveled across time zones were scheduled to participate only after a delay of one week per hour of time difference, to allow for circadian realignment. Eligibility of the volunteers was assessed by telephone call using a general information survey of health and sleep habits. Detailed participant’s characteristics of our final sample are presented in Table 1.

**Table 1.**
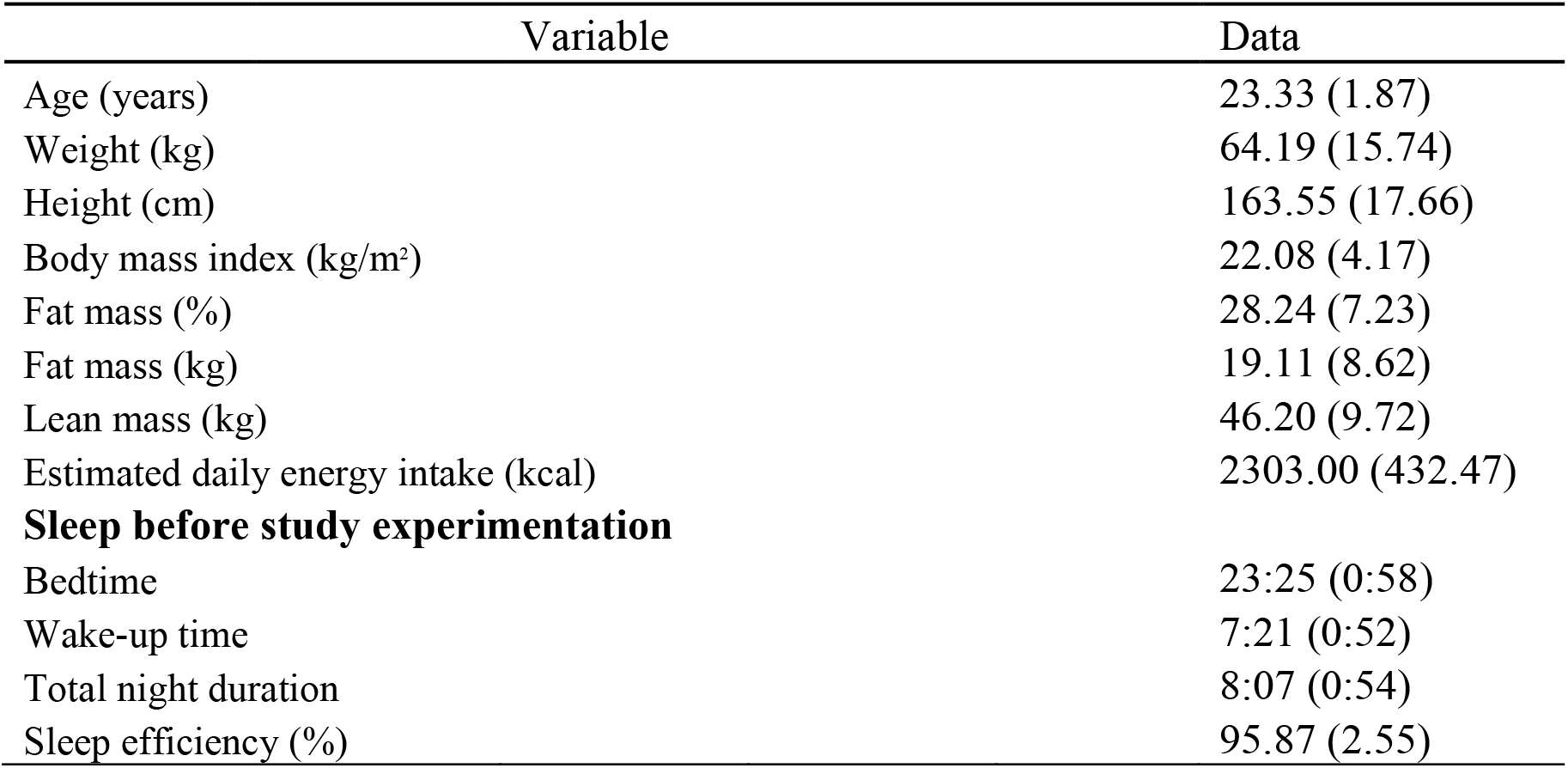
Participant’s characteristics (men, n = 6, women, n = 5). Note. Variables are expressed as mean (standard deviation) or as absolute frequency with percentage in brackets. Total night duration = total sleep time; Sleep efficiency = proportion of sleep duration (without wakefulness) to total night duration. n = 8 for fat and lean mass and for sleep data.

### Experimental protocol

#### Pre-experimental (baseline) session (Figure 1.)

**Figure 1.**
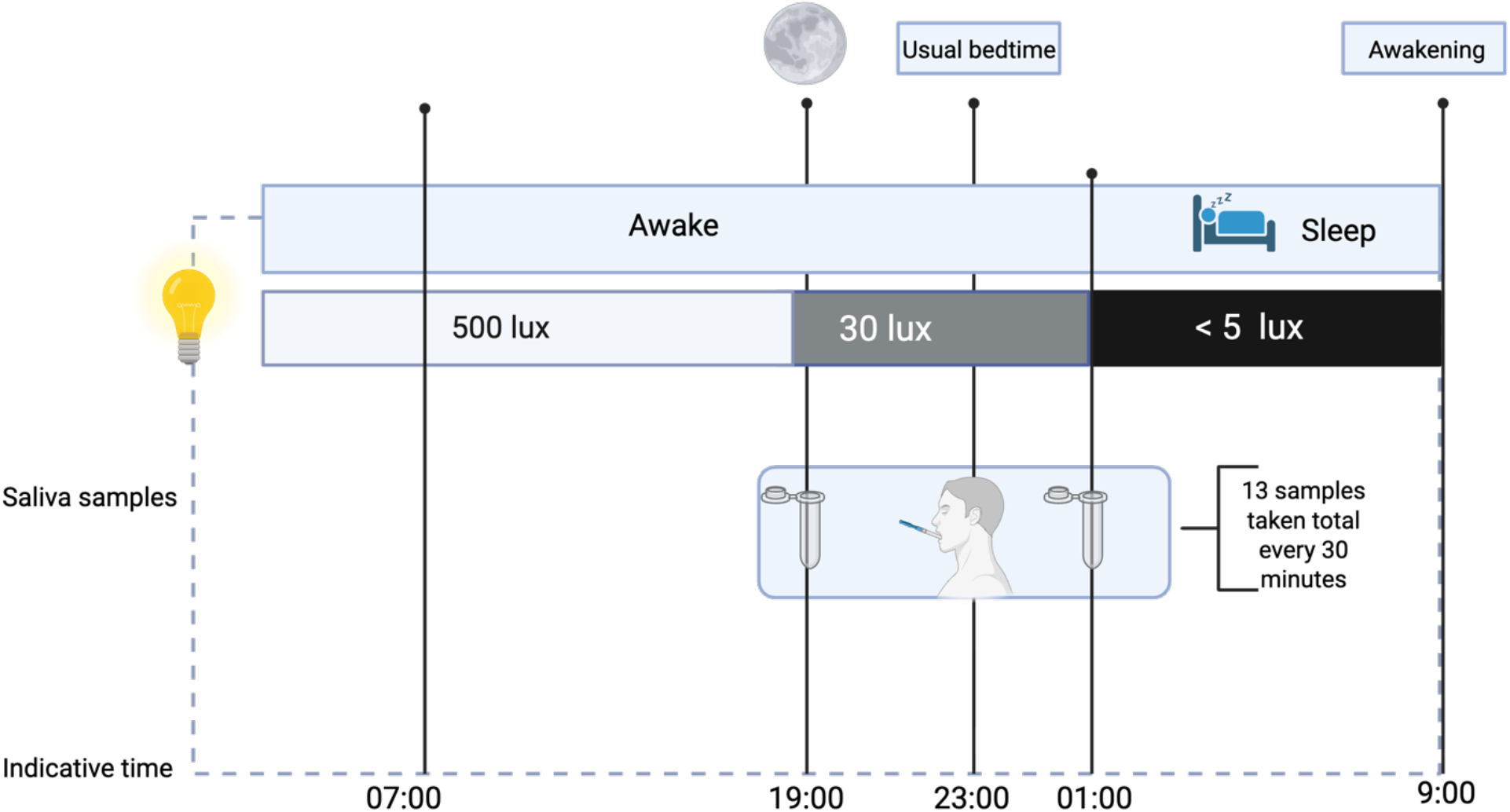
Schematic diagram of pre-experimental (baseline) condition.

Following one week of actigraphy monitoring coupled with sleep diaries to assess baseline sleep-wake patterns, participants arrived at the laboratory approximately 6 hours prior to their habitual bedtime. Upon arrival, an indirect calorimetry measurement was taken to assess their basal metabolic rate, followed by a meal.

Saliva samples were collected beginning 4 hours before the participants’ usual bedtime and continued until 2 hours after. Throughout the sampling period, participants remained mostly seated or reclined in bed under dim lighting (30 lux), with water as the only permitted intake at least 10 minutes before each saliva sample (Pandi-Perumal et al., 2008). Note: Dim lighting (30 lux) was initiated 30 minutes prior to the sampling period to mimic typical sundown patterns. Participants were then allowed to sleep for up to 8 hours in near-total darkness (5 lux) after which they were permitted to shower, have breakfast, and leave the laboratory.

- Indicative times presented in Figures 1,2,and 3 illustrate a representative participant schedule during the testing protocol. As testing was individually aligned with each participant’s habitual sleep-wake patterns, the timing of assessments and interventions varied across individuals, yet adhered to a consistent overall structure.

**Figure 2.**
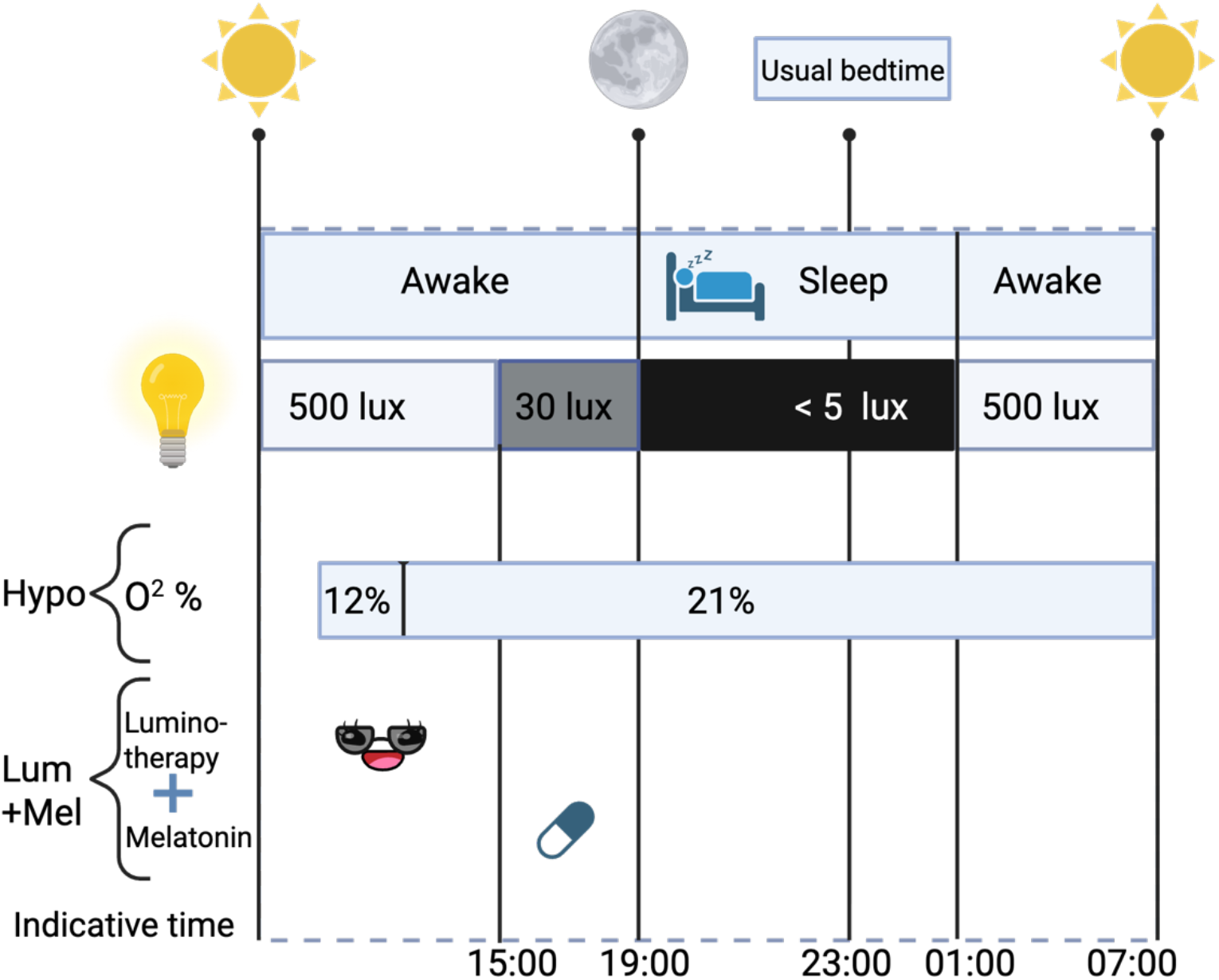
Schematic diagram of experimental conditions (day 1).

**Figure 3.**
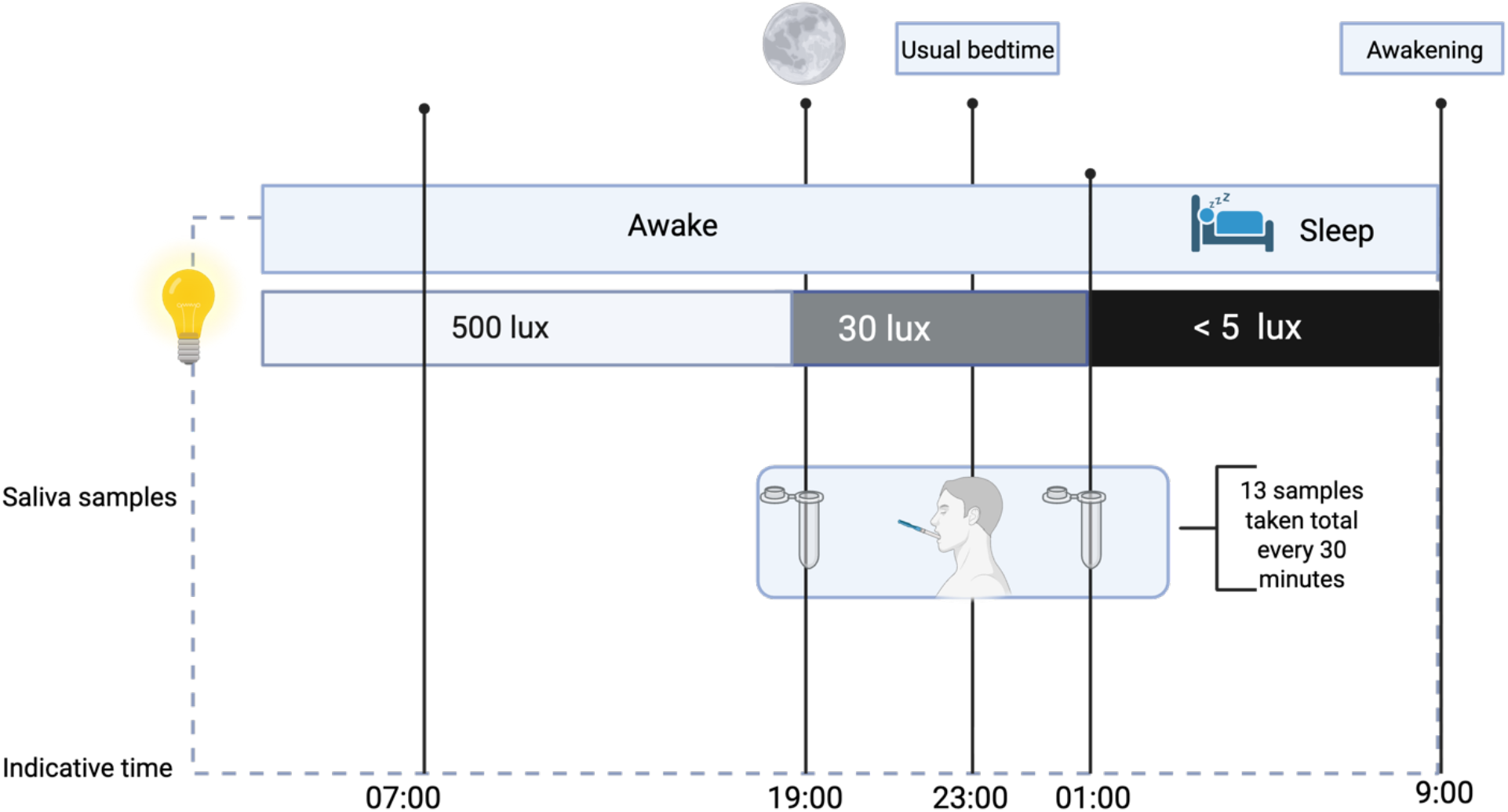
Schematic diagram of experimental conditions (day 2).

#### Experimental Sessions

On the first day of the experimental conditions, participants arrived at the laboratory within an hour of their habitual wake-up time. Upon arrival, they consumed a pre-calculated breakfast (see below; estimation of daily energy expenditure). Two hours after their usual wake-up time, they were exposed to one of two experimental conditions.

- Hypoxia exposure (Hypo): Participants were exposed to normobaric hypoxia (FiO_2_ ∼ 12%) for 2 hours via an oro-nasal mask (see section ‘hypoxic intervention’ justification in discussion). Before and after the hypoxia exposure, participants completed the Lake Louise Scale, a tool for assessing altitude sickness.
- Combined luminotherapy and exogenous melatonin (Lum+Mel): Participants underwent 3 hours of light therapy using Re-Timer glasses and were given an oral dose of 5 mg exogenous melatonin, administered 6 hours before their usual bedtime. During the light therapy, participants could engage in personal activities (e.g., reading, movies, or personal work).

Throughout the day, participants were free to engage in personal activities (no physical activity was allowed) within the laboratory, with no access to external time cues. Of note, participants were not allowed electronic devices with clocks, but had access to a movie filled, clockless laptop during the entire study for entertainment, including during the hypoxic and luminotherapy interventions but excluding the final 30 minutes before bedtime. The lighting was dimmed to 30 lux 4 hours before their usual bedtime. Participants were required to go to bed 4 hours earlier than their usual bedtime in a dark room (<5 lux). They were accorded an 8-hour sleep opportunity, where participants awakened 4 hours prior to their usual wake time.

Upon waking on day 2 of the experimental sessions, the lights were increased to 30 lux, and then to over 500 lux after 20 minutes. Participants were given an hour to shower, get dressed, and have breakfast. The rest of the day mirrored the previous day: they could engage in personal activities (no physical activity), meals were served 4 hours earlier than usual, and the lighting was dimmed to 30 lux 4 hours before their usual sunset time.

In the evening, the same saliva sampling process used during the baseline session was employed. Participants were then allowed to sleep for up to 8 hours, and upon waking. Finally, participants were given time to shower, have breakfast, and leave the laboratory.

### Chronotype assessment

Participants chronotype was assessed using the Morningness-Eveningness Questionnaire (MEQ) developed by (Horne & Östberg, 1976), a widely used 19-question multiple choice questionnaire that evaluate individual preferences for timing of daily activities and sleep-wake patterns. The total score classifies individuals into five chronotype categories: (a) clearly morning (score 16-30), (b) slightly morning (31-41), (c) intermediate (42-58), (d) slightly evening (59-69), and (e) clearly evening (70-86). This questionnaire enabled us to ensure that participants did not have an extreme chronotype, either distinctly morning or distinctly evening. In our sample, the mean MEQ score was 58.9 (SD = 7.9, median = 62.0), indicating a cohort that, on average, fell at the upper end of the intermediate range, bordering on a slightly evening chronotype. These results confirmed that no participants exhibited extreme morning or evening preferences, supporting the inclusion of a relatively homogeneous group in terms of chronotype. Potential participants with extreme chronotypes were excluded from the study. The MEQ is considered a gold standard for chronotype assessment, demonstrating good internal consistency, with a Cronbach’s coefficient of 0.77 to 0.86 (Di Milia et al., 2013).This questionnaire was administered to each participant once during the initial session.

### Estimation of daily energy expenditure

Energy expenditure was measured using indirect calorimetry (VIASYS Healthcare, USA) in a thermoneutral dark room for 30 min at least 4h after food ingestion. Participant was instructed to lie down while oxygen consumption and carbon dioxide production were continuously recorded. The measured energy expenditure was multiplied by a physical activity factor of 1.375 to estimate total daily caloric expenditure corresponding to a low activity level (Harris & Benedict, 1918). This estimate was used to provide participants with an eucaloric diet during each laboratory session to maintain a stable weight and prevent energy deficits that could confound metabolic measurements.

### Sleep assessement

One week before the experimental conditions, participants wore a wrist-worn actigraph (MicroWatch from Ambulatory Monitoring, NY, USA) on their non-dominant wrist to objectively assess sleep-wake patterns. The device records light exposure and motor activity using a linear accelerometer and a miniaturized microprocessor. Participants were instructed to wear the actigraph continuously, removing it only when showering. Actigraphy data were used to determine habitual sleep onset and wake times, total sleep duration, sleep efficiency, and variability in sleep schedules. These metrics ensured that participants maintained stable and consistent sleep patterns prior to the start of laboratory testing. Participants also completed a daily sleep diary each morning of the actigraphy monitoring period. The sleep diary was used to corroborate the data obtained from the actigraphy and to establish the participants’ usual wake-up and bedtimes, which we relied on to develop the tailored laboratory schedule.

### Exposure to normobaric hypoxia

Participants were exposed to normobaric hypoxic air through a tightly fitted oro-nasal mask equipped with a Hans Rudolph non-rebreathing valve, which prevented re-inhalation of expired air. The mask was connected to a system of tubing linked to oxygen extractors (CAT12; Altitude Control Technologies, Lafayette, CO), which were adjusted to provide an inspired oxygen fraction (FiO_2_) of ∼12%, leading to oxyhaemoglobin saturation (SpO_2_) of approximately 80% (Table 2.). The use of SpO_2_ to adjust the hypoxia intensity was chosen as a more individualized approach compared to fixing FiO_2_, which does not account for physiological variability between individuals (Soo et al., 2020). Hypoxic exposure commenced two hours after each participant’s habitual wake-up time. Although we know that sympathetic system activation rises rapidly early in the morning (Lambert et al., 2014), we chose to administer the intervention 2 hours after wake time, which was the earliest possible moment within the constraints of our at-home sleep protocol, to ensure a realistic logistical transition to laboratory measurements. Participants continuously inhaled the hypoxic gas mixture for 120 minutes. Throughout the exposure, heart rate (HR) and SpO_2_ were continuously monitored using a pulse oximeter. Participants remained in a semi-recumbent position in a bed during the hypoxic exposure to prevent any potential falls.

**Table 2.** Physiological responses to normobaric hypoxia. Note. n = 3 for normoxia values; n = 6 for hypoxia values and Lake Louise scale. BPM = beats per minute. The Lac Louise scale ranges from 0 (no symptoms) to 3 (severe symptoms). ^*^ Significant difference (p < .05) compared with values before/after hypoxia (p < .05).

|  | Average (SD) | Min | Max |
| --- | --- | --- | --- |
| **Oxyhaemoglobin saturation (%)** |  |  |  |
| Normoxia | 97.20 (2.25) | 95.8 | 99.8 |
| Hypoxia | 81.22 (2.35)* | 78.25 | 83.79 |
| **Heart rate (BPM)** |  |  |  |
| Normoxia | 85.93 (10.99) | 77 | 98.2 |
| Hypoxia | 92.95 (13.29)* | 79.89 | 115.2 |
| **Lake Louise scale** |  |  |  |
| Headache |  |  |  |
| Before | 0 (0) | 0 | 0 |
| After | 0.7 (0.8) | 0 | 2 |
| Nausea |  |  |  |
| Before | 0 (0) | 0 | 0 |
| After | 0.2 (0.4) | 0 | 1 |
| Weakness or tiredness |  |  |  |
| Before | 0.2 (0.4) | 0 | 1 |
| After | 1.7 (1.0)* | 1 | 3 |
| Vertigo or dizziness |  |  |  |
| Before | 0 (0) | 0 | 0 |
| After | 0.8 (1.2) | 0 | 1 |

### Assessment of hypoxia side-effects

The Lake Louise self-assessment questionnaire was developed to estimate the severity of acute mountain sickness, which refers to the symptoms that may be experienced at high altitudes (Roach et al., 2018). In our study, it was used to document potential symptoms resulting from exposure to reduced oxygen levels. It was administered before and after exposure to hypoxia. The version used included four questions addressing (a) headaches, (b) gastrointestinal symptoms, (c) fatigue/weakness, and (d) dizziness/light-headedness. Participants rated the severity of their symptoms on a 4-point Likert scale, ranging from 0 (no symptoms) to 3 (severe symptoms).

### Oxyhemoglobin saturation

During the hypoxic intervention, participants oxyhemoglobin saturation was monitored using a fingertip pulse oximeter (Masimo Radical 7 unit, Irvine, CA, USA Masimo, USA), which recorded heart rate and peripheral oxygen saturation (SpO_2_) at one-second intervals. The primary objective was to ensure that heart rate remained within normal and SpO_2_ did not fall below 75% during exposure. Under normal oxygen conditions (normoxia), where FiO_2_ is around 21%, SpO_2_ typically ranges from 95% to 98% (Brooks et al., 2005). In our study, SpO_2_ was intentionally maintained around 80% during the 2-hours hypoxic intervention. To confirm the effectiveness of the hypoxic manipulation, a subsample of normoxia data was collected from 3 participants less than 5 minutes before the start of the intervention. Due to technical issues, heart rate and SpO_2_ data for three of the nine participants under hypoxia were not recorded during the experiments. The HR and SpO_2_ values were averaged in one-minute intervals, first individually for each participant and then for the 6 participants whose data were successfully recorded.

### Luminotherapy

Light therapy was administered using Re-Timer light glasses (Re-Timer Pty Ltd, Adelaide, AU), which feature light-emitting diodes (LEDs) positioned on the lower frame. These LEDs emit light at a wavelength of approximately 500 nm, within the short-wavelength (blue-green) spectrum of 435-540 nm, which has been shown to effectively advance the circadian phase (Warman et al., 2003; Wright et al., 2004). The device emits a light intensity of 506 Lux at the eye level, equivalent to standard office lighting. Blue-green light requires lower intensities than traditional white light to achieve similar circadian effects (Warman et al., 2003), and the emitted light is non-blinding and considered safe for ocular exposure. (Burke et al., 2013) demonstrated that a single 3-hour session of light therapy session, either alone or in combination with exogenous melatonin can significantly advance circadian timing. Accordingly, a 3-hour exposure duration session was selected, as longer durations have shown diminishing returns in phase-shifting efficacy (Cheng et al., 2021). To align with participants’ habitual sleep schedules and allow time for laboratory arrival, light therapy began two hours after each participant’s usual wake-up time. This timing has been validated as effective for promoting circadian phase advancement (Janse van Rensburg et al., 2021).

### Exogenous melatonin

Participants received a single 5 mg dose of over-the counter melatonin (Adrien Gagnon, Santé Naturelle A.G. Ltée., Brossard, Qc., CA) administered 6 hours prior to their habitual bedtime. This timing has been identified as optimal for inducing circadian phase advance, and this dose has been shown to be effective when administered once and combined with light therapy (Burke et al., 2013).

### Salivary melatonin

Saliva samples were collected using Salimetrics (College State, PA, USA) salivette swabs. Salivary melatonin levels were measured in duplicates using the Salimetrics competitive immunoassay following the manufacturer’s instructions. All the samples collected from the same participant were run on the same 96-well plate. Variability among duplicates averaged 5.1 ± 6.6 % which is within acceptable limits for salivary melatonin measured using a Salimetrics competitive immunoassay.

### Dim-Light Melatonin Onset calculations

The dim light melatonin onset (DLMO) served as the marker of central clock timing. DLMO was determined using the hockey-stick method which applies a pairwise linear-parabolic function to identify the inflection point marking the transition from baseline to rising melatonin levels (Danilenko et al., 2014). In some sessions (4) the DLMO clearly appeared to occur prior to the 6-hours sampling window. In these cases, the DLMO was estimated using a linear function fitted to the linear portion of the melatonin data. The equation from the fitted line was then used to calculate the time at which it intersects the baseline melatonin values observed in the other sessions of the same participant. Because the normal initial rise in melatonin is curvilinear, the method we used to estimate missing DLMO likely provides values that are “late” compared to the real DLMO. In other words, because the current experimental design involved a phase advance, our method of estimated DLMOs likely correspond to a marginally smaller shift. As such, our method of estimating the missed DLMO should be considered conservative.

### Statistical analyses

A linear mixed model analysis was conducted using Jamovi to determine the effect of experimental condition (i.e., BL, Hypo, and Lum+Mel) on DLMO timing. In the model, participant ID was included as a random intercept to account for within-subject variability. Post hoc comparisons were adjusted using the Bonferroni correction to control for multiple testing.

## Results

### Participant’s characteristics

Table 1 shows the descriptive characteristics of all participants. The final sample consisted of 11 healthy participants (6 men, 5 women, mean age 23 years) characterized by normal adiposity levels. Sleep parameters, assessed using actigraphy over a period of 8 days before the start of the laboratory experiment, confirmed that all individuals met the study’s inclusion criteria.

### Physiological responses to normobaric hypoxia

The physiological effects of the hypoxic intervention are shown in Table 3. Note that the data shown in Table 3 under normoxia are the mean SpO_2_ and HR a few minutes before the start of the intervention for a sub-sample of 3 participants. As expected in healthy individuals, exposure to acute normobaric hypoxia led to a marked drop in oxyhaemoglobin saturation, reflecting reduced oxygen availability. On average, saturation decreased from 97.20% (±2.25) in normoxia to 81.22% (±2.35) in hypoxia—a 16% relative reduction. Heart rate increased significantly, rising from 85.93 bpm (±10.99) under normoxic conditions to 92.95 bpm (±13.29) during hypoxia. Participants reported mild symptoms on the Lake Louise Scale during hypoxia, with average post-exposure scores of 0.7 (±0.8) for headache, 0.2 (±0.4) for nausea, 1.7 (±1.0) for weakness or tiredness, and 0.8 (±1.2) for vertigo or dizziness.

### Salivary melatonin levels

Salivary melatonin levels changed over time (main effect of time: *p* < 0.001), increasing progressively over 6 hours. Compared to BL, Hypo and Lum+Mel led to different average melatonin levels (main effect of condition: all *p* < 0.001). There was no time x condition interaction for salivary melatonin levels (*p* < 0.854.)

### Circadian phase advancement – DLMO

The time course of mean salivary melatonin levels for each experimental condition (BL, Hypo, and Lum+Mel) is illustrated in Figure 5. On average, DLMO occurred 1.30 hour (78 minutes) earlier in the Lum+Mel condition compared to baseline (Tukey p=0.001). In the hypoxia condition, the DLMO occurred on average 0.58 hour (34.8 minutes) earlier than baseline, but this change did not reach statistical significance (Tukey p=0.156).

## Discussion

In this study, we examined whether a 2h exposure to normobaric hypoxia (FiO_2_ ∼ 12%) could promote circadian resynchronization in humans undergoing a phase advance protocol. Specifically, we examined whether acute hypoxia alone could induce circadian phase shifts comparable to those achieved through combined light therapy and exogenous melatonin during a simulated jet lag protocol involving a 4-hour forward shift. To do this, a phase advance protocol was implemented and compared with an established circadian resynchronization treatment – luminotherapy combined with exogenous melatonin intake. Compared to the baseline condition, the combination of luminotherapy and exogenous melatonin produced a statistically significant phase advance in DLMO timing, averaging approximately 78 minutes. After normobaric hypoxia exposure, DLMO occurred 34.8 minutes earlier than baseline; but this shift did not reach statistical significance. Although the magnitude of the phase advance in the hypoxia condition was smaller than that observed with Lum+Mel condition, the difference between the two treatments was not statistically significant, suggesting that both interventions may exert comparable effects on circadian phase under certain condition, albeit with varying degrees in robustness. Our findings reaffirm that luminotherapy combined with exogenous melatonin (Lum+Mel) effectively promotes a phase advance in human circadian rhythms, consistent with previous studies (Burke et al., 2013; Cheng et al., 2021; Pandi-Perumal et al., 2008; Paul et al., 2011). Importantly, our study also reveals that both Hypo and Lum+Mel interventions, when administered alongside a phase-advanced light/dark schedule, are associated with distinct magnitudes of circadian phase shifts. While the absence of a control group undergoing only the light/dark phase advance limits our ability to fully isolate the specific contributions of each intervention, the lack of a statistically significant difference between the Hypo and Lum+Mel conditions suggests that hypoxia may hold potential as a circadian resynchronization strategy. Although the 34.8-minute DLMO advance observed in the hypoxia condition did not reach statistical significance, it may nonetheless hold clinical relevance. Given the small sample size and the possibility that our intervention timing did not optimally align with participants’ individual phase response curve (PRC), it is possible that the circadian system is less sensitive to hypoxia than to light, or that the hypoxic stimulus was not administered at the most effective circadian phase. Future studies are therefore warranted to elucidate the underlying mechanisms, determine optimal timing relative to the PRC, and characterize the dose-response relationship of hypoxic exposure. Despite these limitations, our protocol demonstrates robustness and provides a promising foundation into hypoxia as a novel circadian intervention.

### Hypoxia as a circadian resetting cue: rodent vs. human evidence

Compared to (Adamovich et al., 2017; Post et al., 2025), our findings did not quite replicate the circadian phase-shifting effects observed in previous studies. Unlike Adamovich and colleagues, who assessed circadian phase through locomotor activity over multiple days, our study evaluated circadian markers in the evening the following day of the intervention. This methodological difference may partially account for the absence of a significant phase advance in our results. It is possible that the effects of hypoxia on the human circadian system require a longer duration to manifest. In contrast, Post and colleagues employed a constant posture protocol and exposed human participants to normobaric hypoxia (FiO_2_ ≈ 15%) for approximately 6.5 hours during the early night (Post et al., 2025). The discrepancy between their findings and ours may be explained by several methodological differences, including differences in the timing, duration, and intensity of hypoxia exposure. Additionally, our ability to detect circadian phase shifts may have been challenged by the limited period of salivary sample collection, which may have compromised DLMO estimation. These considerations underscore the importance of future studies with extended monitoring period to better characterize the potential of hypoxia as a circadian phase-shifting intervention. Existing research shows that the effects of luminotherapy and melatonin can be detected as early as the following day—something our study also confirmed (Burke et al., 2013; Paul et al., 2011). It’s also worth noting that simply shifting environmental time cues forward, as we did in our simulated jet lag protocol, can naturally promote circadian realignment at a rate of about one hour per day (Janse van Rensburg et al., 2021). Taken together, our results suggest that while hypoxia remains a promising avenue, more research is needed to understand its timeline and effectiveness compared to established methods like luminotherapy and melatonin.

### Hypoxic intervention

(Adamovich et al., 2017) demonstrated that a 2-hour exposure to 14% fraction of inspired oxygen (FiO_2_) effectively resynchronized the circadian clock in mice following a 6-hour phase advance. Our study used a comparable hypoxic protocol, but with an individualized approach: peripheral capillary oxygen saturation (SpO_2_) was maintained at around 80%. Although Adamovich and colleagues did not report SpO_2_ values, rodent data suggest that FiO_2_ levels of 15% and 12% yield average SpO_2_ values of 81.4% and 72.9%, respectively (Morgan et al., 2014). It is therefore reasonable to infer that the 14% FiO_2_ used by Adamovich and colleagues resulted SpO_2_ levels like those observed in our participants (81.22 ± 2.35%). Thus, our hypoxic intervention appears like that performed in mice in the study by Adamovich and colleagues and has the potential to have induced the expected physiological effects in humans. This suggests that the lack of significant difference between the hypoxic and BL conditions is unlikely to result from a difference in hypoxemic level of our hypoxic intervention. However, interspecies differences in hypoxia sensitivity remain a critical consideration. Moreover, Adamovich and colleague’s administered hypoxia during the mice’s active (dark) phase, whereas our participants remained at rest in a semi-recumbent position during the early part of their active (daytime) phase. While increased physical activity in mice could have lowered their SpO_2_ levels, this effect could have been offset by their greater pulmonary gas exchange efficiency compared to humans (Gonzalez & Kuwahira, 2018). The timing of the hypoxic intervention in our protocol was guided by Adamovich and colleagues protocol and adapted for human circadian rhythms (Adamovich et al., 2017). Specifically, we initiated hypoxia approximately two hours after habitual wake time, coinciding with the phase-advance portion of the human PRC to light. This alignment allowed for a direct comparison between hypoxia and light exposure. The timing was therefore chosen so that the hypoxia and luminotherapy sessions would start at the same time, enabling a close comparison of their effects. Timing is a well-established determinant of efficacy in circadian interventions, including light and melatonin (Burgess & Emens, 2016) and it is possible that hypoxia follows a phase response curve. However, no such curve has yet been established. A recent study in mice (Manella et al., 2020) showed that the timing of hypoxia exposure —whether during the light or dark phase, —alters the expression of clock genes, with responses varying across tissues. These effects appear to be driven primarily by the circadian clocks themselves, rather than by the light-dark cycle. Notably, a 4-hour exposure to 6% O_2_ during the light phase advanced the phase in the lungs and kidneys but delayed it in the liver. These findings suggest that hypoxia can shift circadian rhythms, but its effects may differ across peripheral clocks and depend on timing. However, only one circadian time point was tested, limiting conclusions about temporal dynamics (Manella et al., 2020). In humans, evidence remains limited.

### Luminotherapy and exogenous melatonin intake intervention

Luminotherapy and exogenous melatonin administration are well-established methods for circadian phase shifting and were used as a reference intervention in our study to compare against the effects of hypoxia. Our results showed a significant difference in DLMO between the combined Lum+Mel condition and BL, but not between hypoxia and the other conditions. This indicates that the Lum+Mel intervention effectively advanced the circadian phase relative to baseline, though not significantly more than hypoxia. These results reinforce the evidence that even a single combined treatment of luminotherapy and exogenous melatonin is effective for advancing the circadian phase (Cheng et al., 2021). In our study, luminotherapy was administered two hours after participants’ usual wake time. Research shows that luminotherapy can effectively shift circadian phase when given within a window extending six hours before or after the core body temperature nadir (CBTn), with peak effectiveness observed between 0 to 3 hours from the CBTn (Janse van Rensburg et al., 2021). In our protocol, the luminotherapy treatment occurred approximately 4 hours after the CBTn—later than in the studies by (Burke et al., 2013) and (Paul et al., 2011)—however, still yielding a significant effectiveness. This consolidates Janse van Rensburg et al., 2021)’s effective 6-hour window. Accordingly, although the timing of light exposure was later than in some other studies, we observed a measurable effect of our (Lum+Mel) leading to a significant average DLMO phase advance of 78 minutes compared to our BL condition.

As for exogenous melatonin, we administered a 5 mg dose six hours before participants’ usual bedtime, following the protocol established by (Burke et al., 2013). The timing of melatonin administration is known to influence its efficacy, with higher doses generally producing greater phase shifts when taken earlier relative to DLMO, while lower doses are more effective when administered closer to DLMO (Burgess et al., 2010). For example, (Burgess et al., 2008) reported that a 3 mg dose produced the largest phase advance when taken five hours before DLMO. Based on this, we expected similar outcomes to those seen in (Burke et al., 2013), which our findings indeed confirmed. Specifically, our Lum+Mel condition produced a DLMO phase advance of 0.59 hours relative to baseline—consistent with the expected range of 0.62 to 1.13 hours reported in previous studies involving similar protocols (Burke et al., 2013; Paul et al., 2011). This range also accounts for the additive effect of our jet lag protocol, which independently contributes to circadian phase advancement. It is worth noting that studies have found significant variability in the actual melatonin content of over-the-counter supplements, raising concerns about whether the administered dose truly equaled 5 mg (Grigg & Ianakieva, n.d.). This variability, along with individual difference in circadian responsiveness, may account for the range of responses observed. Our average phase advance of approximately 1.3 hours falls within, or slightly above, the expected range of 0.62 to 1.13 hours reported in previous studies— particularly given the inclusion of a jetlag protocol, which alone is known to promote circadian phase advances.

### Endogenous salivary melatonin as a circadian marker

As previously reported, we did not observe a statistically significant difference in melatonin levels between the Hypo and BL, which limits our ability to draw firm conclusions about the role of hypoxia as a circadian resynchronizer. Similarly, there was no significant difference in DLMO timing between Hypo and Lum+Mel, suggesting that Lum+Mel did not produce a significantly greater phase-shifting effect than hypoxia alone. Nevertheless, we believe that in some participants, we may have narrowly missed capturing the exact timing of their DLMO. Indeed, several participants exhibited salivary melatonin concentrations above our predetermined DLMO threshold of 3 pg/mL at the start of sampling, suggesting that their DLMO may have occurred just prior to our sampling window. This raises the possibility that a circadian phase advance occurred but went undetected due to the restricted sampling period. Unfortunately, the limited number of salivary melatonin samples available prevents us from confirming or ruling out this interpretation.

In addition, salivary melatonin levels in the present study exhibited the expected circadian profile, increasing progressively over time across the 6-h sampling period (main effect of time: all *p* < 0.001) (Figure 4.). Notably, melatonin levels also differed significantly by experimental condition (main effect of condition: all *p* < 0.001), though no significant condition by time interaction was observed (*p* = 0.854), suggesting that while baseline levels were modulated by condition, the overall temporal dynamics of melatonin levels were not significantly different between Hypo and Lum+Mel. Melatonin levels were significantly higher under Lum+Mel, then Hypo followed with BL condition having the lowest quantity levels of melatonin. This could possibly be explained by increased sympathetic activation following hypoxia (Post et al., 2025) and/or a priming effect of exogenous melatonin intake.

**Figure 4.**
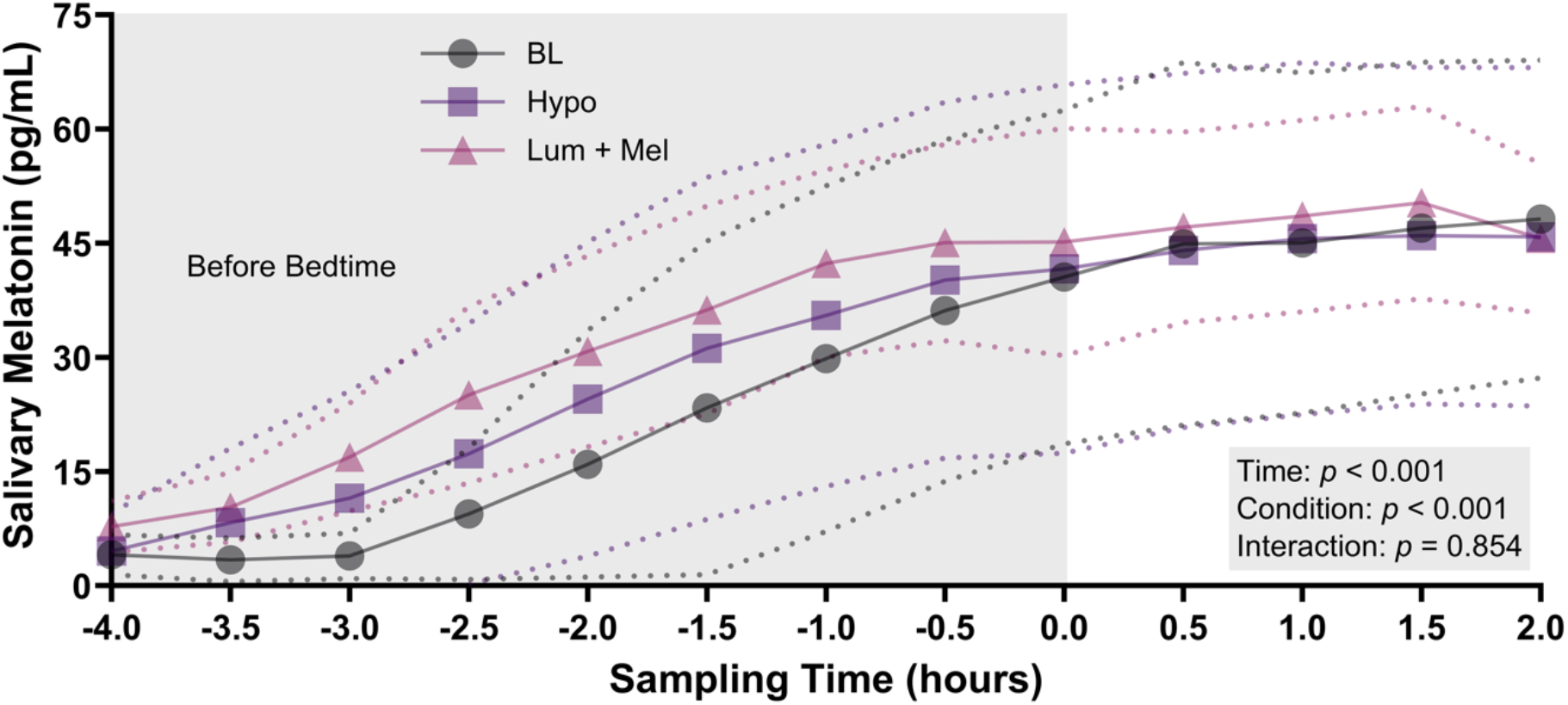
Mean salivary melatonin levels measured in samples collected every 30 minutes over a 6-hour period, beginning 4 hours before and ending 2 hours after participants’ habitual bedtime. Data are presented for a baseline condition and two experimental conditions. Melatonin levels are plotted against sampling time, expressed in hours relative to habitual bedtime (−4.0 to +2.0). Values are shown as means (solid lines) with 95% confidence intervals (dotted lines).

**Figure 5.**
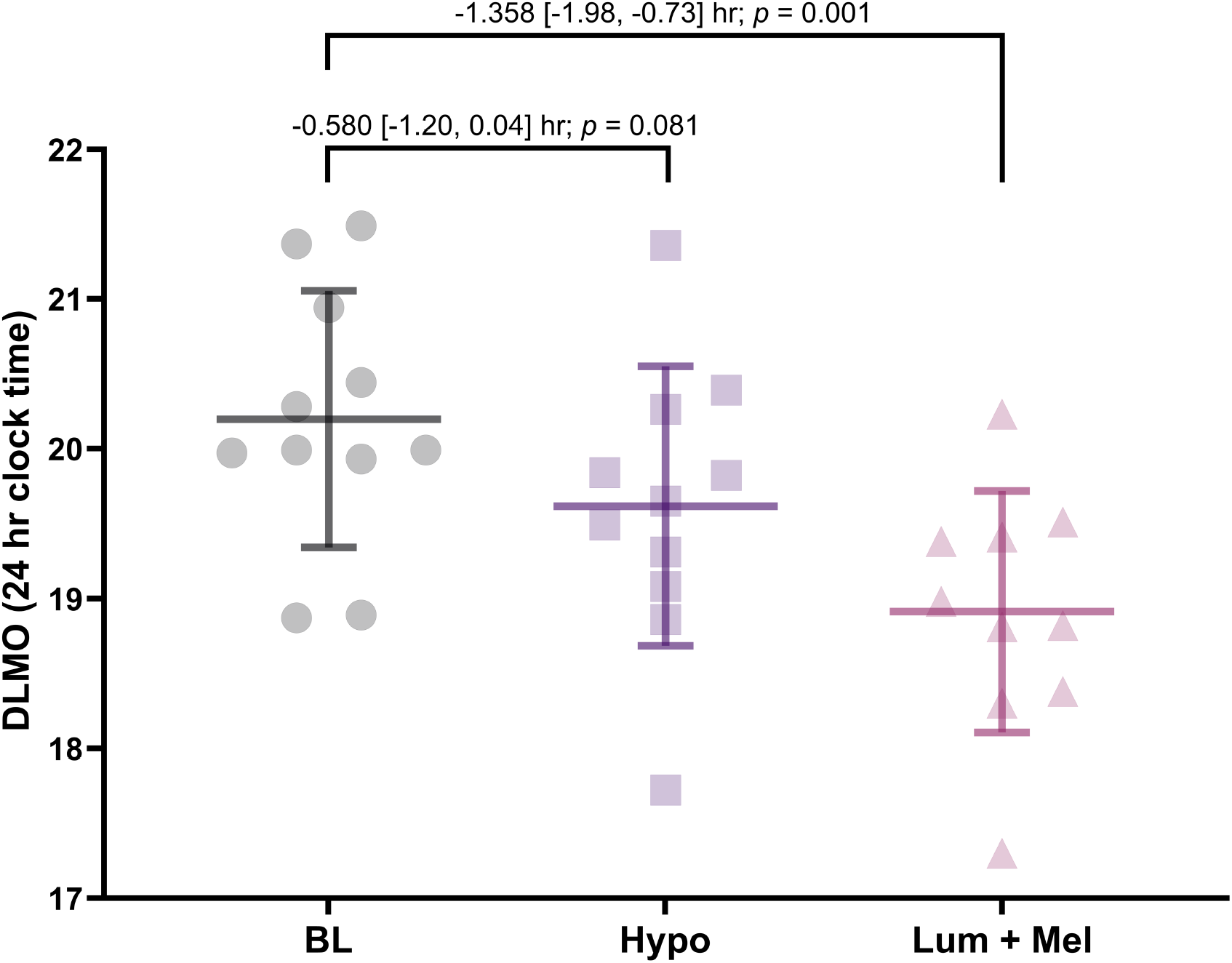
Dim Light Melatonin Onset (DLMO) times across experimental conditions (Baseline [BL], Hypoxia [Hypo], and Light + Melatonin [Lum+Mel]), calculated using the hockey stick method (Danilenko et al., 2014). DLMO is expressed in 24-hour clock time (17:00–22:00) to reflect the actual time of melatonin onset. Data are presented as group means (horizontal bars) with standard deviations (vertical brackets) and individual participant values (dots). Between-condition differences are shown as mean [95% CI].

### Masking effect of our experimental design

To accurately assess circadian markers like endogenous melatonin, it is crucial to minimize external influences (e.g., light, activity, food intake) that can mask internal rhythms. Masking refers to the direct, clock-independent effects of environmental factors on behavior, distinct from entrainment, which involves synchronization of the internal circadian clock (Mrosovsky, 1999). The constant routine (CR) protocol was developed to isolate endogenous rhythms by controlling environmental and behavioral variables (Rietveld & Waterhouse, n.d.). CR requires participants to remain awake in a semi-recumbent position under constant conditions for at least 24 hours, with evenly spaced meals and minimal activity (Broussard et al., 2017; Martinez-Nicolas et al., 2013). However, CR is resource-intensive, induces sleep deprivation, and is unsuitable for repeated measures (Broussard et al., 2017; Kenny et al., 1996; Lack & Lushington, 1996; Martinez-Nicolas et al., 2013). Due to these limitations and participant burden, we did not implement a CR protocol. As such, masking effects—particularly from light exposure—remain present in our baseline condition, which was conducted under natural environmental conditions (see Figure 1). Our Hypo and Lum+Mel conditions introduced a 4-hour phase advance in light exposure to simulate jet lag and assess the additive effects of light therapy, melatonin, and hypoxia. While this design allowed us to test our interventions, it limits our ability to fully isolate their effects on DLMO phase shifts due to unavoidable masking.

### Strength and weaknesses of the study

This exploratory study employed a within-subject randomized crossover design, in which each participant experienced all experimental conditions. This approach enabled direct comparisons of each participant’s responses against their own baseline, thereby minimizing inter-individual variability and enhancing the statistical robustness of our findings. To mitigate potential order effects, the sequence of experimental conditions was counterbalanced, with the baseline condition always conducted first to allow for an adaptation night in the laboratory. Furthermore, to account for differences in circadian timing, the experimental protocol was carefully personalized so that circadian interventions were administered at each participant’s optimal time.

This study also has several non negligeable limitations that hinder potential conclusions to be drawn from our data. First, there is a selection bias as most of our participants were healthy university students thereby limiting the generalizability of our findings. While strict inclusion criteria were necessary to reduce confounding variables, they further narrowed the representativeness of the sample. The final sample size was also very limited due to data loss, recruitment difficulties, the demanding nature of the protocol, and the significant time commitment (totaling nearly 115 hours in the lab) required from the participants. A larger sample would strengthen the external validity of conclusions regarding the effects of hypoxia. Additionally, while participants had free access to a clockless laptop for entertainment, its use was restricted during the 30 minutes preceding bedtime to minimize potential effects on sleep onset. Although the device provided no time cues, its unmonitored use during the day could represent a minor uncontrolled factor potentially influencing circadian responses. Another limit lies in the use of a single circadian marker, salivary melatonin. This marker is also susceptible to masking effects from light, food intake, and sleep, which are challenging to fully control a jet lag protocol. Nonetheless, our study protocol was specifically designed to tightly control these variables, thereby reducing their potential confounding effects. Also, the baseline condition (BL) was subject to different masking influences compared to the experimental conditions (Hypo and Lum+Mel). Creating a true constant routine protocol to establish a more accurate baseline would have required an extended 48-hour period without environmental changes and still may not have resolved the issue of differential masking. Additionally, the absence of a phase shift protocol using a placebo intervention is a limitation, as it would have allowed us to isolate the specific effect of hypoxia on salivary melatonin. This decision was based on the inherent difficulty of simulating a placebo for hypoxia. Instead, we chose to compare hypoxia with a well-established intervention: light therapy combined with exogenous melatonin. The use of over-the-counter exogenous melatonin tablets is another limitation of this study, as variability in melatonin concentration may be present (Grigg & Ianakieva, n.d.).

## Conclusion

Circadian desynchronization affects a considerable number of people worldwide and is associated with significant short and long term physical and mental health consequences. Addressing this issue requires both the refinement of existing interventions and the development of novel, more effective strategies. In this exploratory study, we investigated the potential of hypoxia as a circadian resynchronization intervention. However, the methodological limitations we identified prevent us from drawing definitive conclusions about hypoxia’s effectiveness in this context. Despite this, our study lays the groundwork for more in-depth future research on the effects of hypoxia on the human circadian system. Importantly, our study was able to confirm findings from previous research by demonstrating that the reference intervention – luminotherapy combined with exogenous melatonin – was effective in resynchronizing the circadian clock following a phase advance protocol. By highlighting the limitations of our exploratory work and proposing directions for future studies, we aim to guide ongoing research in this field, which is vital for human health. Evidence from rodent studies and new findings supports the potential of hypoxia to reset circadian rhythms, offering a promising avenue to help millions of people restore a healthy biological rhythm.

## Acknowledgements

We would also like to extend our sincere thanks to all the daytime, evening, and nighttime supervisors who supported the smooth running of the experimental sessions, as well as to Professor Stewart Foguel for his valuable and constructive feedback on the protocol. RM is a recipient of a PhD’s scholarship from the Natural Sciences and Engineering Research Council of Canada. Pascal Imbeault is a recipient of a research chair from the Institut du Savoir Montfort (2016-018-Chair-PIMB) and a Discovery Grant from the Natural Sciences and Engineering Research Council of Canada (NSERC) (RGPIN-2019-792 04438).

## References

Adamovich, Y., Ladeuix, B., Golik, M., Koeners, M. P., & Asher, G. (2017). Rhythmic Oxygen Levels Reset Circadian Clocks through HIF1α. Cell Metabolism, 25(1), 93–101. 10.1016/j.cmet.2016.09.014

Bin, Y. S., Postnova, S., & Cistulli, P. A. (2019). What works for jetlag? A systematic review of non-pharmacological interventions. Sleep Medicine Reviews, 43, 47–59. 10.1016/j.smrv.2018.09.005

Brooks, G., Fahey, T., & Baldwin, K. (2005). Exercise Physiology: Human Bioenergetics and Its Application.

Broussard, J. L., Reynolds, A. C., Depner, C. M., Ferguson, S. A., Dawson, D., & Wright, K. P. (2017). Circadian rhythms versus daily patterns in human physiology and behavior. In V. Kumar (Ed.), Biological Timekeeping (pp. 279–295). Springer. 10.1007/978-81-322-3688-7_13

Burgess, H. J., & Emens, J. S. (2016). Circadian-Based Therapies for Circadian Rhythm Sleep-Wake Disorders. Current Sleep Medicine Reports, 2(3), 158–165. 10.1007/s40675-016-0052-1

Burgess, H. J., Revell, V. L., & Eastman, C. I. (2008). A three pulse phase response curve to three milligrams of melatonin in humans. The Journal of Physiology, 586(2), 639–647. 10.1113/jphysiol.2007.143180

Burgess, H. J., Revell, V. L., Molina, T. A., & Eastman, C. I. (2010). Human Phase Response Curves to Three Days of Daily Melatonin: 0.5 mg Versus 3.0 mg. The Journal of Clinical Endocrinology & Metabolism, 95(7), 3325–3331. 10.1210/jc.2009-2590

Burke, T. M., Markwald, R. R., Chinoy, E. D., Snider, J. A., Bessman, S. C., Jung, C. M., & Wright, K. P. (2013). Combination of Light and Melatonin Time Cues for Phase Advancing the Human Circadian Clock. Sleep, 36(11), 1617–1624. 10.5665/sleep.3110

Cheng, D. C. Y., Ganner, J. L., Gordon, C. J., Phillips, C. L., Grunstein, R. R., & Comas, M. (2021). The efficacy of combined bright light and melatonin therapies on sleep and circadian outcomes: A systematic review. Sleep Medicine Reviews, 58, 101491. 10.1016/j.smrv.2021.101491

Coste, O., Beaumont, M., Batéjat, D., Beers, P. V., Charbuy, H., & Touitou, Y. (2004). Hypoxic depression of melatonin secretion after simulated long duration flights in man. Journal of Pineal Research, 37(1), 1–10. 10.1111/j.1600-079X.2004.00128.x

Coste, O., Beaumont Maurice, Batéjat, Denise, Van Beers, Pascal, & and Touitou, Y. (2004). Prolonged Mild Hypoxia Modifies Human Circadian Core Body Temperature and may be Associated with Sleep Disturbances. Chronobiology International, 21(3), 419–433. 10.1081/CBI-120038611

Coste, O., Van Beers, P., & Touitou, Y. (2009). Hypoxia-induced changes in recovery sleep, core body temperature, urinary 6-sulphatoxymelatonin and free cortisol after a simulated long-duration flight. Journal of Sleep Research, 18(4), 454–465. 10.1111/j.1365-2869.2009.00744.x

Coste, O., Van Beers, Pascal, & and Touitou, Y. (2007). Impact of Hypobaric Hypoxia in Pressurized Cabins of Simulated Long-Distance Flights on the 24 h Patterns of Biological Variables, Fatigue, and Clinical Status. Chronobiology International, 24(6), 1139–1157. 10.1080/07420520701800702

Crowley, S. J., & Eastman, C. I. (2012). Melatonin in the Afternoons of a Gradually Advancing Sleep Schedule Enhances the Circadian Rhythm Phase Advance. Psychopharmacology, 225(4), 825. 10.1007/s00213-012-2869-8

Danilenko, K. V., Verevkin, E. G., Antyufeev, V. S., Wirz-Justice, A., & Cajochen, C. (2014). The hockey-stick method to estimate evening dim light melatonin onset (DLMO) in humans. Chronobiology International, 31(3), 349–355. 10.3109/07420528.2013.855226

Di Milia, L., Adan, A., Natale, V., & Randler, C. (2013). Reviewing the psychometric properties of contemporary circadian typology measures. Chronobiology International, 30(10), 1261–1271. 10.3109/07420528.2013.817415

Gonzalez, N. C., & Kuwahira, I. (2018). Systemic Oxygen Transport with Rest, Exercise, and Hypoxia: A Comparison of Humans, Rats, and Mice. Comprehensive Physiology, 8(4), 1537–1573. 10.1002/j.2040-4603.2018.tb00050.x

Grigg, -Damberger Madeleine M., & Ianakieva, D. (n.d.). Poor Quality Control of Over-the-Counter Melatonin: What They Say Is Often Not What You Get. Journal of Clinical Sleep Medicine, 13(02), 163–165. 10.5664/jcsm.6434

Harris, J. A., & Benedict, F. G. (1918). A Biometric Study of Human Basal Metabolism. Proceedings of the National Academy of Sciences, 4(12), 370–373. 10.1073/pnas.4.12.370

Hofstra, W. A., & de Weerd, A. W. (2008). How to assess circadian rhythm in humans: A review of literature. Epilepsy & Behavior, 13(3), 438–444. 10.1016/j.yebeh.2008.06.002

Horne, J. A., & Östberg, O. (1976). A self-assessment questionnaire to determine morningness-eveningness in human circadian rhythms. International Journal of Chronobiology, 4, 97– 110.

Janse van Rensburg, D.C., Jansen van Rensburg, A.x, Fowler, P. M., Bender, A. M., Stevens, D., Sullivan, K. O., Fullagar, H. H. K., Alonso, J.-M., Biggins, M., Claassen-Smithers, A., Collins, R., Dohi, M., Driller, M. W., Dunican, I. C., Gupta, L., Halson, S. L., Lastella, M., Miles, K. H., Nedelec, M., … Botha, T. (2021). Managing Travel Fatigue and Jet Lag in Athletes: A Review and Consensus Statement. Sports Medicine, 51(10), 2029–2050. 10.1007/s40279-021-01502-0

Kenny, G. P., Giesbrecht, G. G., Thoden, J. S., & Kenny, G. (1996). Post-exercise thermal homeostasis as a function of changes in pre-exercise core temperature. European Journal of Applied Physiology and Occupational Physiology, 74(3), 258–263. 10.1007/BF00377448

Kuhlman, S. J., Craig, L. M., & Duffy, J. F. (2018). Introduction to Chronobiology. Cold Spring Harbor Perspectives in Biology, 10(9), a033613. 10.1101/cshperspect.a033613

Lack, L., & Lushington, K. (1996). The rhythms of human sleep propensity and core body temperature. Journal of Sleep Research, 5(1), 1–11. 10.1046/j.1365-2869.1996.00005.x

Lambert, E. A., Chatzivlastou, K., Schlaich, M., Lambert, G., & Head, G. A. (2014). Morning Surge in Blood Pressure Is Associated With Reactivity of the Sympathetic Nervous System. American Journal of Hypertension, 27(6), 783–792. 10.1093/ajh/hpt273

Lewis, P., Korf, H. W., Kuffer, L., Groß, J. V., & Erren, T. C. (2018). Exercise time cues (zeitgebers) for human circadian systems can foster health and improve performance: A systematic review. BMJ Open Sport — Exercise Medicine, 4(1), e000443. 10.1136/bmjsem-2018-000443

Manella, G., Aviram, R., Bolshette, N., Muvkadi, S., Golik, M., Smith, D. F., & Asher, G. (2020). Hypoxia induces a time- and tissue-specific response that elicits intertissue circadian clock misalignment. Proceedings of the National Academy of Sciences, 117(1), 779–786. 10.1073/pnas.1914112117

Martinez-Nicolas, A., Ortiz-Tudela, E., Rol, M. A., & Madrid, J. A. (2013). Uncovering Different Masking Factors on Wrist Skin Temperature Rhythm in Free-Living Subjects. PLOS ONE, 8(4), e61142. 10.1371/journal.pone.0061142

Mistlberger, R. E., & Rusak, B. (2005). Circadian rhythms in mammals: Formal properties and environmental influences. In Principles and practice of sleep medicine (pp. 321–334). Elsevier.

Molina, T. A., & and Burgess, H. J. (2011). Calculating the Dim Light Melatonin Onset: The Impact of Threshold and Sampling Rate. Chronobiology International, 28(8), 714–718. 10.3109/07420528.2011.597531

Morgan, B. J., Adrian, R., Bates, M. L., Dopp, J. M., & Dempsey, J. A. (2014). Quantifying hypoxia-induced chemoreceptor sensitivity in the awake rodent. Journal of Applied Physiology, 117(7), 816–824. 10.1152/japplphysiol.00484.2014

Mrosovsky, N. (1999). Masking: History, Definitions, and Measurement. Chronobiology International, 16(4), 415–429. 10.3109/07420529908998717

Pandi-Perumal, S. R., Trakht, I., Spence, D. W., Srinivasan, V., Dagan, Y., & Cardinali, D. P. (2008). The roles of melatonin and light in the pathophysiology and treatment of circadian rhythm sleep disorders. Nature Clinical Practice Neurology, 4(8), 436–447. 10.1038/ncpneuro0847

Paul, M. A., Gray, G. W., Lieberman, H. R., Love, R. J., Miller, J. C., Trouborst, M., & Arendt, J. (2011). Phase advance with separate and combined melatonin and light treatment. Psychopharmacology, 214(2), 515–523. 10.1007/s00213-010-2059-5

Post, T. E., De Gioannis, R., Schmitz, J., Wittkowski, M., Schäper, T. M., Wrobeln, A., Fandrey, J., Schmitz, M.-T., Takahashi, J. S., Jordan, J., Elmenhorst, E.-M., & Aeschbach, D. (2025). Resetting of the Human Circadian Melatonin Rhythm by Ambient Hypoxia. Journal of Pineal Research, 77(1), e70029. 10.1111/jpi.70029

Reid, K. J., & Abbott, S. M. (2015). Jet Lag and Shift Work Disorder. Sleep Medicine Clinics, 10(4), 523–535. 10.1016/j.jsmc.2015.08.006

Rietveld, W. J., & Waterhouse, J. M. (n.d.). Circadian Rhythms and Masking: An Overview.

Roach, R. C., Hackett, P. H., Oelz, O., Bärtsch, P., Luks, A. M., MacInnis, M. J., & Baillie, J. K. (2018). The 2018 Lake Louise Acute Mountain Sickness Score. High Altitude Medicine & Biology, 19(1), 4–6. 10.1089/ham.2017.0164

Soo, J., Girard, O., Ihsan, M., & Fairchild, T. (2020). The Use of the SpO2 to FiO2 Ratio to Individualize the Hypoxic Dose in Sport Science, Exercise, and Health Settings. Frontiers in Physiology, 11. 10.3389/fphys.2020.570472

Warman, V. L., Dijk, D.-J., Warman, G. R., Arendt, J., & Skene, D. J. (2003). Phase advancing human circadian rhythms with short wavelength light. Neuroscience Letters, 342(1), 37– 40. 10.1016/S0304-3940(03)00223-4

Weinert, D., & Waterhouse, J. (2017). Interpreting Circadian Rhythms. In Biological Timekeeping: Clocks, Rhythms and Behaviour (pp. 23–45). Springer, New Delhi. 10.1007/978-81-322-3688-7_2

Wright, H. R., Lack, L. C., & Kennaway, D. J. (2004). Differential effects of light wavelength in phase advancing the melatonin rhythm. Journal of Pineal Research, 36(2), 140–144. 10.1046/j.1600-079X.2003.00108.x

